# refMLST: Reference-based Multilocus Sequence Typing Enables Universal Bacterial Typing

**DOI:** 10.1101/2023.06.12.544669

**Authors:** Mondher Khdhiri, Ella Thomas, Chanel de Smet, Priyanka Chandar, Vivek Madasu, Induja Chandrakumar, Jean M Davidson, Paul Anderson, Samuel D Chorlton

## Abstract

**Summary:** Commonly used approaches for genomic investigation of bacterial outbreaks, including SNP and gene-by-gene approaches, are limited by the requirement for curated allele schemes. As a result, they only work on a select subset of known organisms, and fail on novel or less studied pathogens. We introduce refMLST, a gene-by-gene approach using the reference genome of a bacterium to form a scalable, reproducible and robust method to perform outbreak investigation. When applied to 1263 *Salmonella enterica* genomes, refMLST enabled consistent clustering, improved resolution and faster processing in comparison to chewieSnake. refMLST is applicable to any bacterial species with a public genome, does not require a curated scheme, and automatically accounts for genetic recombination.

**Availability and Implementation:** refMLST is freely available for academic use at https://bugseq.com/academic.

## Introduction

Traditionally, approaches for detecting and investigating bacterial outbreaks using DNA sequencing are divided into single-nucleotide polymorphism (SNP) and gene-by-gene approaches, the latter comprising core genome multilocus sequence typing (cgMLST) and whole genome multilocus sequence typing (wgMLST). Multiple comparisons have demonstrated that these approaches largely perform equivalently, with different strengths and weaknesses [1–3]. SNP-based approaches are negatively impacted by genetic recombination, where a single evolutionary event may introduce many SNPs. Popular methods for recombination correction, such as ClonalFrameML and Gubbins give different outputs for different sets of inputs, and therefore cannot be applied sample-by-sample in real-time investigation of a growing outbreak [4, 5]. Furthermore, popular SNP-based approaches only examine nucleotide sites shared across all isolates; as the outbreak grows, fewer sites are included in analysis and resolution paradoxically decreases [6, 7]. Conversely, gene-by-gene approaches are limited by the need for a stable, curated, centrally-hosted scheme, limiting their utility to select previously-studied pathogens [8–10]. Even with a species-specific scheme, most scheme servers are subject to licensing (Enterobase, cgMLST.org), precluding their use in applied settings. Both approaches may be hampered by computational requirements and speed of analysis; chewieSnake, a recently published cgMLST analysis, took more than a day when analysing thousands of genomes on 8 CPUs, precluding application to large datasets [11].

Recently, hashing of locus sequences has been introduced for gene-by-gene approaches to enable robust reproducibility without the need for curating every previously seen allele of a locus in the centrally-hosted scheme [11, 12]. Instead of naming alleles of a locus in the order in which they were discovered, hash-based methods use information theory techniques to identify an allele based on the allele’s sequence. However, these approaches still require a centrally-hosted scheme to define loci locations for hashing, and do not address other aforementioned issues with SNP and gene-by-gene approaches.

To date, no tool has combined the universal applicability of a reference genome with gene-by-gene analysis, nor enabled recombination-corrected, species-agnostic bacterial outbreak investigation. We present reference-based MSLT (refMLST), a reproducible gene-by-gene approach to type any bacterium based on a single reference genome.

### Theory and Implementation

Below, we detail the implementation of refMLST for identifying reference loci, typing them in a query genome, and combining typing data across query genomes to identify outbreak clusters. An overview of this process is provided in Figure 1. The query genome for refMLST is a genomic assembly in FASTA format which may be generated from whole genome or metagenomic sequencing; however, refMLST requires that the genome is complete and free from contamination (eg. if a mixed isolate was sequenced, imperfect taxonomic binning, or strain duplication). refMLST automatically runs BUSCO on query genomes to assess completeness and contamination, and will only process those with greater than 90% unfragmented single copy orthologs in accordance with MIMAG standards [14, 15]. All other genomes are flagged for review.

**Figure 1.**
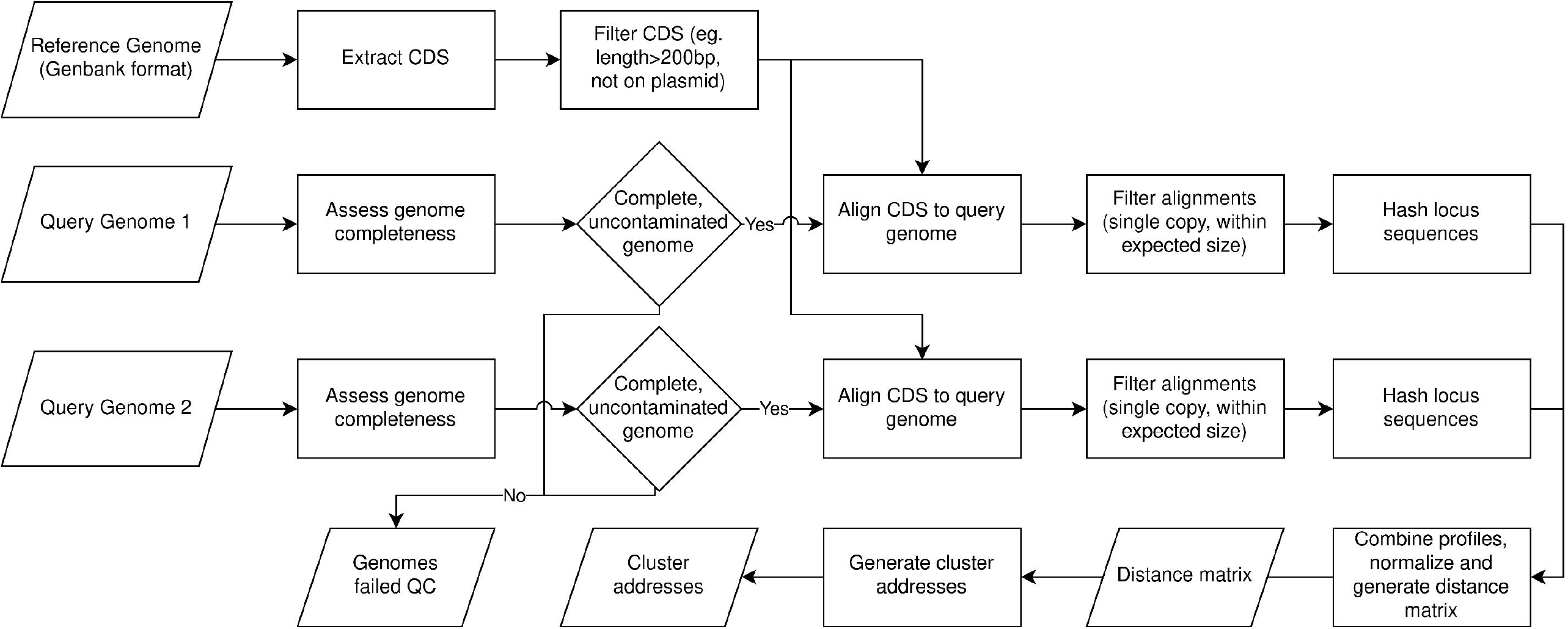
Overview of refMLST workflow. Parallelograms reflect inputs/outputs, rectangles reflect processes and diamonds reflect decision points.

Unlike multilocus sequence typing, cgMLST and wgMLST, refMLST does not require a curated scheme of alleles – it only requires a single reference genome for the pathogen under study, facilitating analysis of any bacterial species. The reference genome in Genbank format is used to perform locus (CDS) identification and extraction from the query genome; this file may be sourced from NCBI for any publicly available genome. For novel pathogens, this file can be annotated from a sequenced reference (eg. any sample from outbreak) using standard genome annotation pipelines. Querying individual loci ameliorates reference bias associated with gene synteny; further steps to ameliorate reference bias are discussed below. While reference genomes of closely related species could be used with refMLST if no reference is available for the same species, we limit our analysis below to the use of reference genomes within the same species, preserving the number of CDS shared between query and reference genomes.

Loci on mobile genetic elements may have a different evolutionary history from the bacterial chromosome, comprise a significant component of the accessory genome, contribute to reference bias, and ultimately confound analysis; they are therefore excluded from analysis based on annotation in the Genbank file. Specifically, any sequence with annotation “plasmid”, “phage”, “virus” or “mobile_element” is excluded. As outbreaks of antimicrobial resistance or other important phenotypes may be caused by plasmids, users are encouraged to combine the results of refMLST for host bacterial typing with dedicated plasmid typing tools for a holistic investigation of outbreaks [16]. Additionally, refMLST omits loci less than 200 base pairs in length in accordance with other gene-by-gene analysis tools [17].

To call alleles in the assembled genome of interest, locus sequences from above are rapidly aligned against the query genome assembly using minimap2 with the following command [18]:

′ ′ ′

minimap2 -a --MD -N 1 --end-bonus 5 query_assembly.fna cds_sequences.fna

′ ′ ′

Minimap2 can align nucleic acid sequences with up to 20% sequence divergence, enabling identification of genes beyond the expected species boundary of 95% absolute nucleotide identity [18, 19]. We opted not to use protein aligners such as DIAMOND or MMSeqs2 as the former requires the query assembly to be translated into amino acid sequence, while the latter cannot account for frameshifts as of Release 14-7e284 [20, 21]. Alignments of loci against the genome assembly are filtered for multiple hits to exclude paralogous loci. To ensure quality locus alignments, reference loci must align end to end, and newly discovered alleles must be within 20% of the size of the reference allele. Next, a CRC32 hash is taken of the allele sequence in the query genome to convert it into a compact, easy to understand number.

Evolutionary distances between isolates are calculated with a hamming distance, ignoring missing alleles. Resulting hamming distances are normalized by the number of loci found in both genomes relative to the reference genome, with the formula

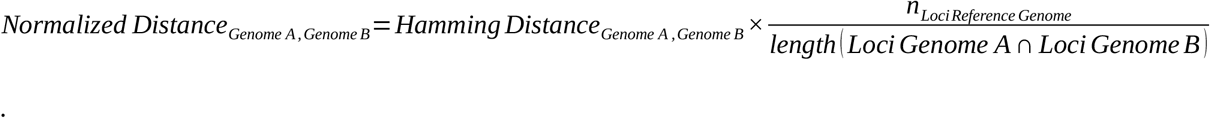

This normalization corrects for imbalances in the presence of reference accessory genes across clades of the analysed organism. The aforementioned genome quality control ensures that small inter-genome distances are not a result of missing loci in either genome when performing pairwise comparisons. refMLST automatically accounts for recombination by examining whole genes instead of individual SNPs to calculate the hamming distance, obviating the need for post-analysis, non-deterministic recombination correction.

Cluster addresses are calculated by iterating through query genomes in the order at which they were sequenced/submitted, performing single-linkage hierarchical clustering. Reproducible cluster addresses reflecting inter-sample allele distances are generated; description of similar cluster addresses has previously been published [11, 22]. In brief, cluster addresses are composed of seven digits, where each digit reflects a distance threshold (1000, 200, 100, 50, 20, 10, 5 alleles). Addresses are read from left to right, and genomes with greater genomic similarity will share more cluster address digits in common. An example of cluster addresses and their utility is provided in Table 1. Unlike other address generation mechanisms, and because typing of each genome is deterministic with refMLST, cluster addresses are also deterministic and stable, and therefore do not need matching across real-time analyses as an outbreak grows [11, 23]. All samples are output in a distance matrix and line list with cluster codes.

**Table 1.** Example of refMLST cluster address output. The reference genome is always assigned the first cluster, 1.1.1.1.1.1.1. Sample 1 is over 1000 alleles from the reference genome, as the first digit (1000 allele address) is different between these two cluster addresses. Sample 2 is more than 10 but less than 20 alleles different from Sample 1, as the 10 allele address is different. Sample 3 is more than 5 but less than 10 alleles different from Sample 2, and more than 10 but less than 20 alleles different from Sample 1. Cluster addresses do not change over time as genomes are added to the outbreak analysis.

| Sample Name | Merged Cluster Address | 1000 Allele Address | 200 Allele Address | 100 Allele Address | 50 Allele Address | 20 Allele Address | 10 Allele Address | 5 Allele Address |
| --- | --- | --- | --- | --- | --- | --- | --- | --- |
| Reference Genome | 1.1.1.1.1.1.1 | 1 | 1 | 1 | 1 | 1 | 1 | 1 |
| Sample 1 | 2.1.1.1.1.1.1 | 2 | 1 | 1 | 1 | 1 | 1 | 1 |
| Sample 2 | 2.1.1.1.1.2.1 | 2 | 1 | 1 | 1 | 1 | 2 | 1 |
| Sample 3 | 2.1.1.1.1.2.2 | 2 | 1 | 1 | 1 | 1 | 2 | 2 |

### Application

refMLST does not require a curated scheme of alleles – it only requires a single reference genome, facilitating analysis of any bacterial species. refMLST automatically accounts for recombination by examining whole genes instead of individual SNPs, obviating the need for post-analysis, non-deterministic recombination correction. Consequently, cluster addresses are deterministic and stable, and therefore do not need matching across real-time analyses as an outbreak grows [11, 23]. We applied refMLST to 1263 *Salmonella enterica* isolates with publicly available sequencing data, and compared results to chewieSnake (v3.0.0), a recently published, hash-based cgMLST tool [11]. We find the overall correlation coefficient between refMLST and chewieSnake allele distances to be 1.0 (Figure 2). At distances less than 100 refMLST alleles, refMLST distances are on average 20 (SD: 10) alleles greater than chewieSnake, indicating an enhanced ability to resolve closely related isolates. Increased refMLT allele distances are explained by examination of a greater number of loci when comparing genomes (refMLST: 4271 loci, chewieSnake: 3000 loci). Accounting for differences in allele distances between tools with linear regression, we find outbreak clusters at comparable allele distances (refMLST: 20 alleles, chewieSnake: 10 alleles) to be highly correlated with an Adjusted Rand Index of 0.92 [24]. refMLST is also rapid: refMLST processed the 1263 *S. enterica* genomes in 59 minutes on an r6i.2xl EC2 instance with 8 CPUs and 64 GB of RAM; in contrast, chewieSnake took 23 hours and 33 minutes with the same data and server.

**Figure 2.**
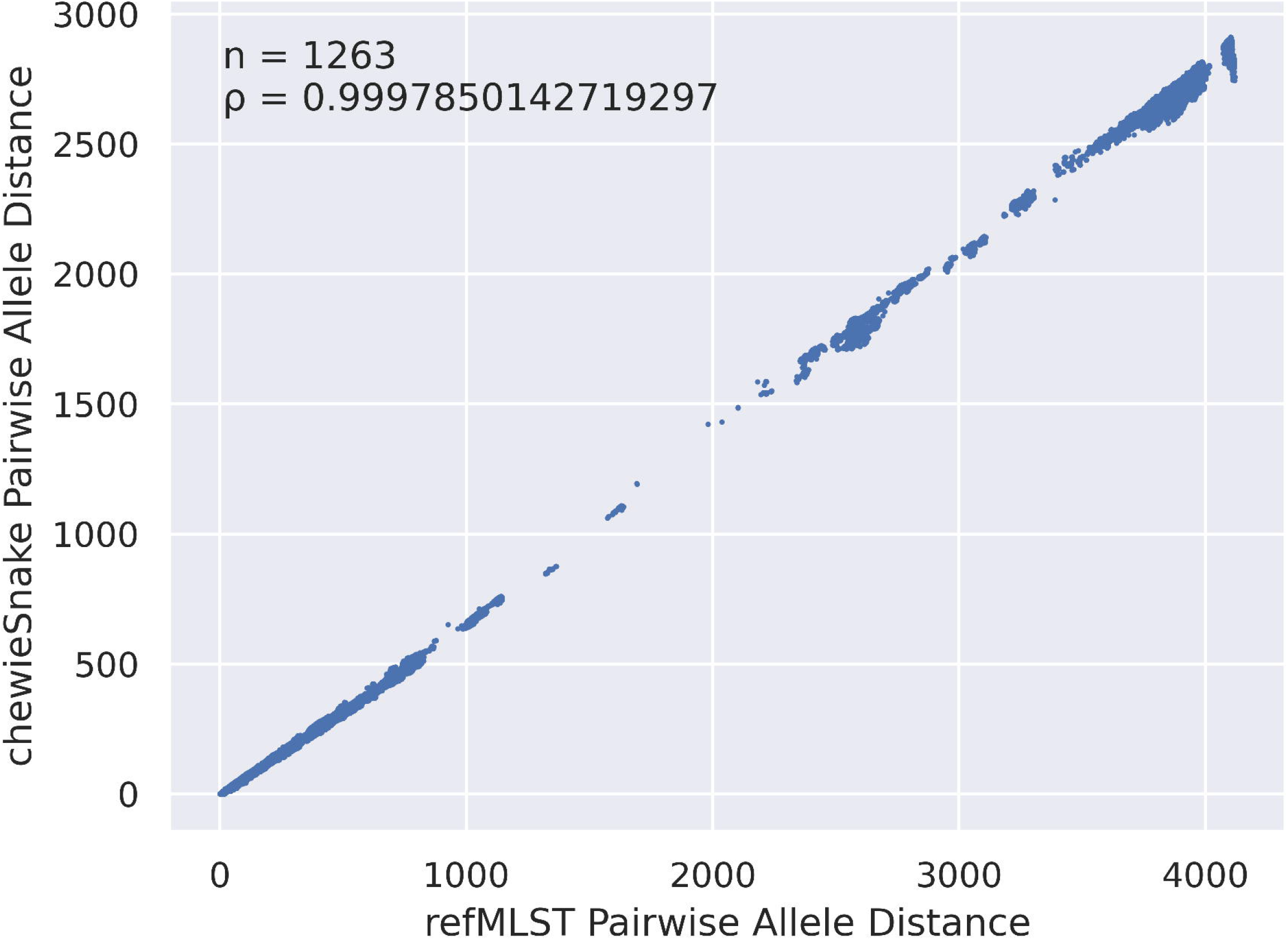
refMLST pairwise allele distances compared with chewieSnake pairwise allele distances for 1263 *Salmonella enterica* genomes. Each dot represents a comparison of two genomes.

## Conclusion

refMLST combines the advantages of SNP and gene-by-gene approaches to enable reproducible, universal bacterial typing without suffering from each approach’s limitations.

## Data Summary

*Salmonella enterica* genomes are available from [11] (https://doi.org/10.5281/zenodo.4338293). refMLST is freely available for academic use at https://bugseq.com/academic. Documentation on refMLST, including input requirements and interpretation of outputs is available at https://docs.bugseq.com.

## Acknowledgements

The authors are grateful to the clinical and public health laboratories who have evaluated, validated and provided feedback on the routine use of refMLST for outbreak investigation.

## Funding

The author(s) received no financial support for the research, authorship, and/or publication of this article.

## Conflict of Interest

IC and MK are employees of BugSeq Bioinformatics Inc. SDC is a shareholder in BugSeq Bioinformatics Inc.

## Notes

https://doi.org/10.5281/zenodo.4338293

### Competing Interest Statement

IC, MK and SDC have been employed by BugSeq Bioinformatics Inc.

